# A Deep Learning Approach for NeoAG-Specific Prediction Considering Both HLA-Peptide Binding and Immunogenicity: Finding Neoantigens to Making T-Cell Products More Personal

**DOI:** 10.1101/2021.12.22.473942

**Authors:** Xian-Xian Liu, Gloria Li, Wei Luo, Juntao Gao, Simon Fong

## Abstract

**Background:** An emerging type of cancer treatment, known as cell immunotherapy, is gaining popularity over chemotherapy or other radiation therapy that causes mass destruction to our body. One favourable approach in cell immunotherapy is the use of neoantigens as targets that help our body immune system identify the cancer cells from healthy cells. Neoantigens, which are non-autologous proteins with individual specificity, are generated by non-synonymous mutations in the tumor cell genome. Owing to its strong immunogenicity and lack of expression in normal tissues, it is now an important target for tumor immunotherapy. Neoantigens are some form of special protein fragments excreted as a by-product on the surface of cancer cells during the DNA mutation at the tumour. In cancer immunotherapies, certain neoantigens which exist only on cancer cells elicit our white blood cells (body’s defender, anti-cancer T-cell) responses that fight the cancer cells while leaving healthy cells alone. Personalized cancer vaccines therefore can be designed de novo for each individual patient, when the specific neoantigens are found to be relevant to his/her tumour. The vaccine which is usually coded in synthetic long peptides, RNA or DNA representing the neoantigens trigger an immune response in the body to destroy the cancer cells (tumour). The specific neoantigens can be found by a complex process of biopsy and genome sequencing. Alternatively, modern technologies nowadays tap on AI to predict the right neoantigen candidates using algorithms. However, determining the binding and non-binding of neoantigens on T-cell receptors (TCR) is a challenging computational task due to its very large search space.

**Objective:** To enhance the efficiency and accuracy of traditional deep learning tools, for serving the same purpose of finding potential responsiveness to immunotherapy through correctly predicted neoantigens. It is known that deep learning is possible to explore which novel neoantigens bind to T-cell receptors and which ones don’t. The exploration may be technically expensive and time-consuming since deep learning is an inherently computational method. one can use putative neoantigen peptide sequences to guide personalized cancer vaccines design.

**Methods:** These models all proceed through complex feature engineering, including feature extraction, dimension reduction and so on. In this study, we derived 4 features to facilitate prediction and classification of 4 HLA-peptide binding namely AAC and DC from the global sequence, and the LAAC and LDC from the local sequence information. Based on the patterns of sequence formation, a nested structure of bidirectional long-short term memory neural network called local information module is used to extract context-based features around every residue. Another bilstm network layer called global information module is introduced above local information module layer to integrate context-based features of all residues in the same HLA-peptide binding chain, thereby involving inter-residue relationships in the training process. introduced

**Results:** Finally, a more effective model is obtained by fusing the above two modules and 4 features matric, the method performs significantly better than previous prediction schemes, whose overall r-square increased to 0.0125 and 0.1064 on train and increased to 0.0782 and 0.2926 on test datasets. The RMSE for our proposed models trained decreased to approximately 0.0745 and 1.1034, respectively, and decreased to 0.6712 and 1.6506 on test dataset.

**Conclusion:** Our work has been actively refining a machine-learning model to improve neoantigen identification and predictions with the determinants for Neoantigen identification. The final experimental results show that our method is more effective than existing methods for predicting peptide types, which can help laboratory researchers to identify the type of novel HLA-peptide binding.

## 1 Introduction

As major histocompatibility complexes in humans[1], the human leukocyte antigens (HLAs) have important functions to present antigen peptides onto T-cell receptors for immunological recognition and responses. Interpreting and predicting HLA–peptide binding are important to study T-cell epitopes, immune reactions, and the mechanisms of adverse drug reactions. HLAs play an important role in the immune system to present peptides to TCRs for immune responses[2]; however, this process may result in adverse outcomes under certain circumstances. Though various methods and tools have been developed to predict HLA–peptide binding, challenges in this field remain for researchers to address. Most machine learning methods take only peptide sequences as input features; therefore, individual models have to be developed for each HLA. HLA-I proteins are primarily encoded by three genes (HLA-A, HLA-B, and HLA-C), which are widely expressed in most cell types in human. In addition, specialized cell types can express HLA-E, HLA-F, or HLA-G genes. HLA-A, -B, and -C genes (hereafter referred to as HLA-I) are the most polymorphic genes in the human genome and over 12,000 distinct alleles are documented in the human population. Humans have in general different combinations of HLA-I alleles and, therefore, express up to six different HLA-I proteins (two for each gene). HLA-I molecules bind short peptides, mainly 9–11 amino acids, and different HLA-I alleles have distinct binding specificities, which implies that a broad spectrum of peptides can be displayed across different individuals.

In the past several years, a variety of deep learning models such as deep neural network [3], convolutional neural network [4] and recurrent neural network [5] have been developed for advancing HLA-peptide prediction over traditional machine learning algorithms [6]. Meanwhile, some studies weighted greatly on the data quality for improving prediction accuracy. Zhang et al used physicochemical properties to group amino acids in their HMM.49 POPI used 20 physicochemical properties to represent each amino acid according to AAindex database 9.0.58,67 SVRMHC utilized both sparse encoding and their 11-factor physicochemical property descriptors in their method.51,68 They compared both types of descriptors and found that the two types of descriptors showed different performance on different HLA alleles. However, the existing methods have different kinds of limitations such as lack of supported HLAs, a problem dealing with peptides of various lengths and high false positives in experimental validations. We have proposed the network method and a combination approach that uses multiple types of methods to address some of these challenges.

Overcoming this limitation requires developing novel computational methods to model and predicting neopeptide-HLA. A new method for Identifying neo-peptides of HLA. Our aim is to describe the main steps of antigen presentation that proved to be successful in making quantitative predictions of antigens. The more biological aspects of antigen presentation and processing are covered in many other reviews.

In this study, we demonstrate how to prepare the high-quality input for feature extraction including global peptides sequence descriptors and local peptides sequence descriptors for compositions of amino acid properties. We review different types of machine learning methods and tools that have been used for HLA–peptide binding prediction. We introduce a framework that any HLA-peptides binding sequence to a sequence of vector embeddings. We will predict antigen presentation---what could we learn from a million peptides and compare the prediction performances of several neural networks time series forecasting models namely GRU, BiLSTM and our proposed method with existing predictors on the 4 types of novel dataset and We also summarize the descriptors based on which the HLA–peptide binding prediction models have been constructed and discuss the limitation and challenges of the current methods. Lastly, we give a future perspective on the HLA–peptide binding prediction method based on network analysis.

## 2 Data collection

We retrieved proteins from the IEDB automatic server benchmark database at http://tools.iedb.org/auto_bench/mhci/weekly/.that provided HLA-peptide binding sequence data and annotations. In this study, we worked on data sets from public databases. HLA class I alleles that are of HLA-A, B, and C subtypes, we trained our model on 10-length peptides binding to HLA-A, HLA-B and HLA-C alleles with available HLA sequences. Totally, the training dataset contains 121,787 peptide-HLA binding peptides covering 2 HLA-A alleles (9196 samples in HLA-A01:01 and samples in HLA-A03:01), 1 HLA-B alleles (46915 samples) and 1 HLA-C alleles (9196 samples). The detailed information of the training and testing data are listed in Supplementary File. In this work, we only trained and tested on peptides rang form 4 to10-length. We downloaded all available benchmark datasets from and evaluated our model on this dataset.

## 3 Peptides Data representation approach

In this study, we used 4 different feature encoding schemes to encode the peptide sequences into feature vector with fixed length. Feature extraction method converts character representation of a protein sequence into a numerical representation, which uses the corresponding feature vector to represent peptides sequence information.

As depicted in Fig. 1. each input is an HLA-peptide pair. For both the peptide and the HLA in an input, based on amino acid classification and physicochemical properties, the property of amino acids and sequence information are exploited to generate features, i.e., amino acid composition (AAC), the local amino acid composition (LAAC) and dipeptide composition (DC), as well as the local dipeptide composition (LDC)feature. Schematic illustrate the process of physicochemical properties generation for a single input Sequences. Selection and preparation of the samples are as primary outputs for the terminel outputs of machine learning.

**Fig. 1.**
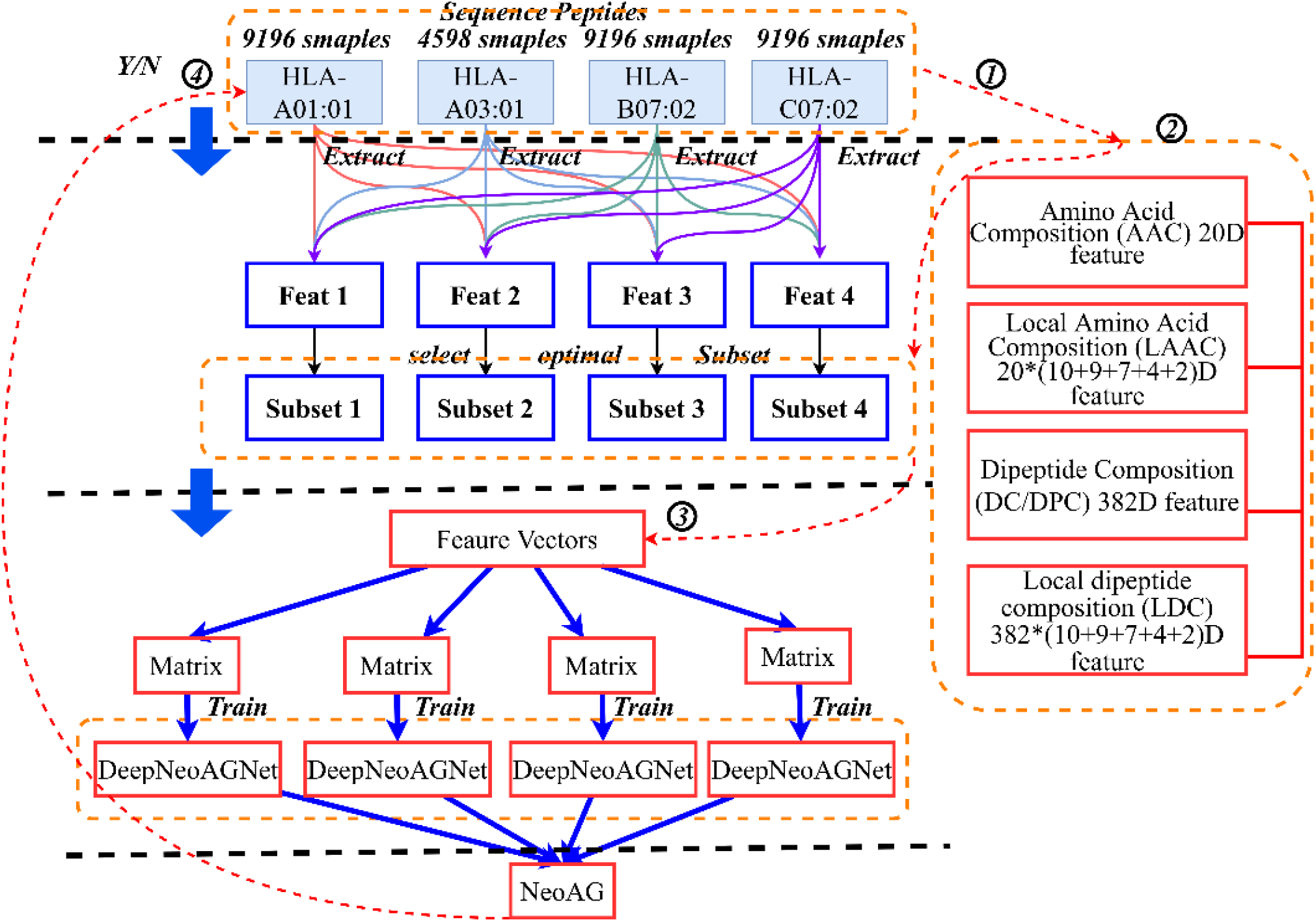
Overview of Main Strategies for the Identification of Neoantigens

**Fig. 2.**
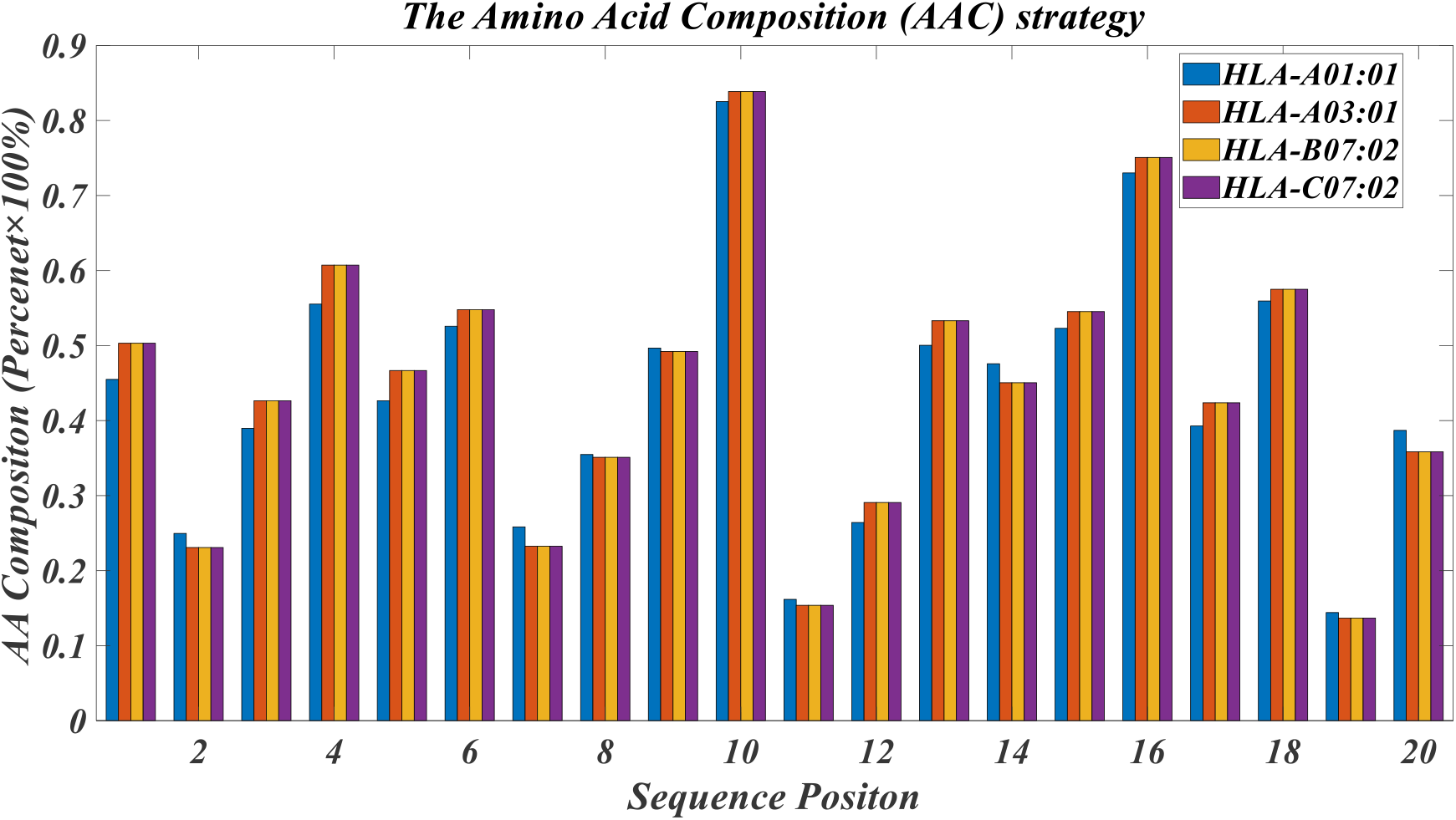
Overall comparison of amino acid composition

**Fig. 3.**
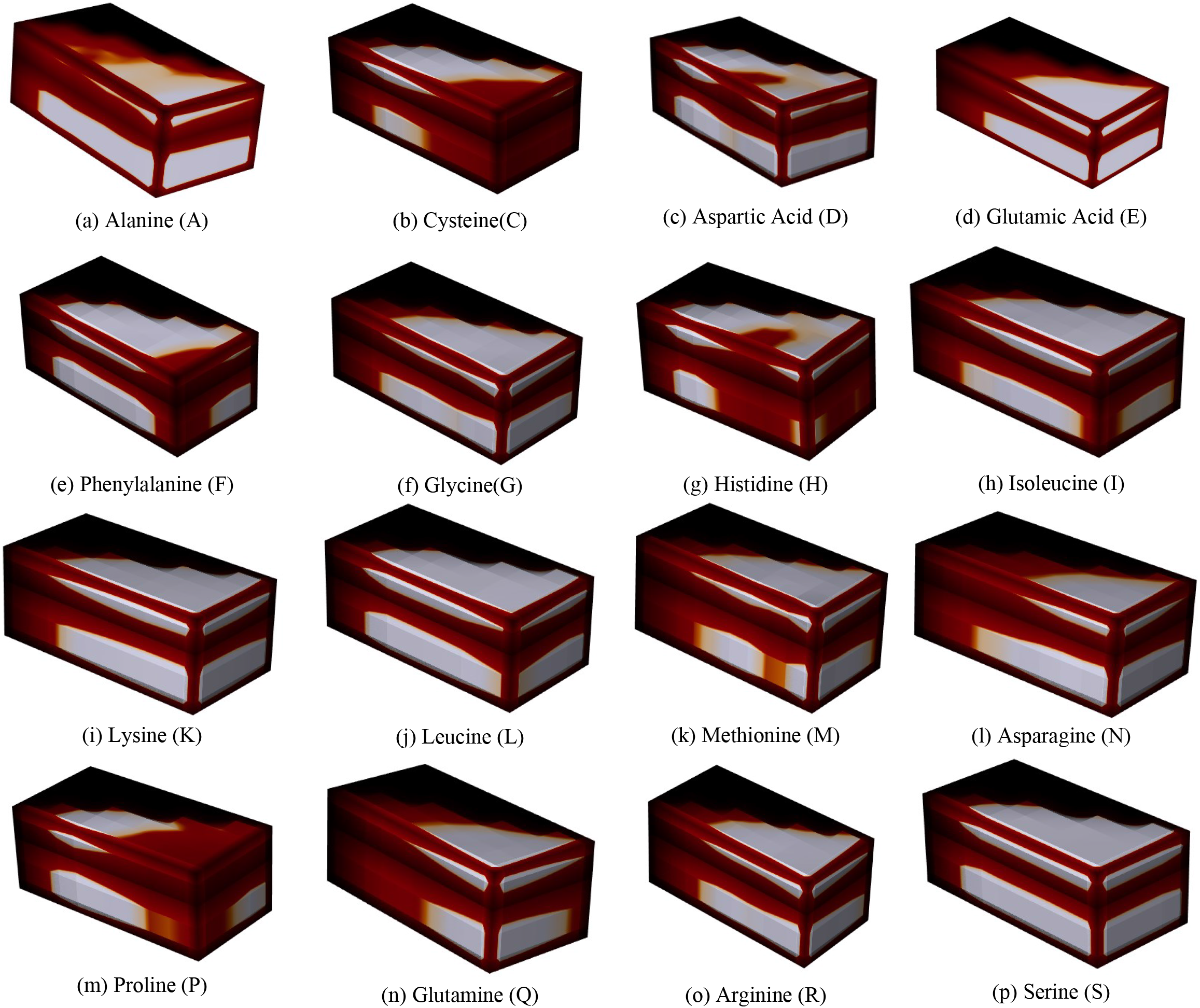
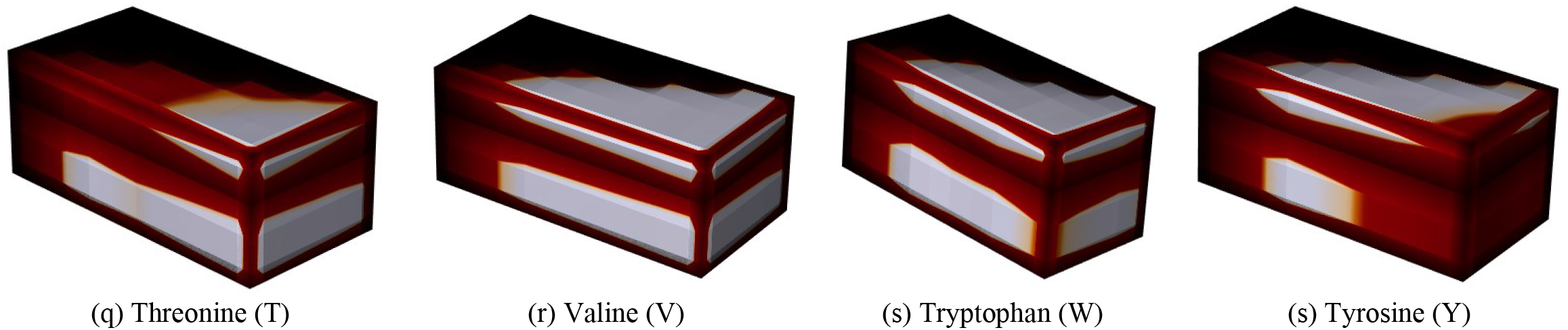
local composition values for each amino acid and window size combination for each peptide. Recall at amino acid level. short side: window size from 10%AA∼80%AA; long y axis: height z axis: the percent of each local amino acid composition.

**Fig. 4.**
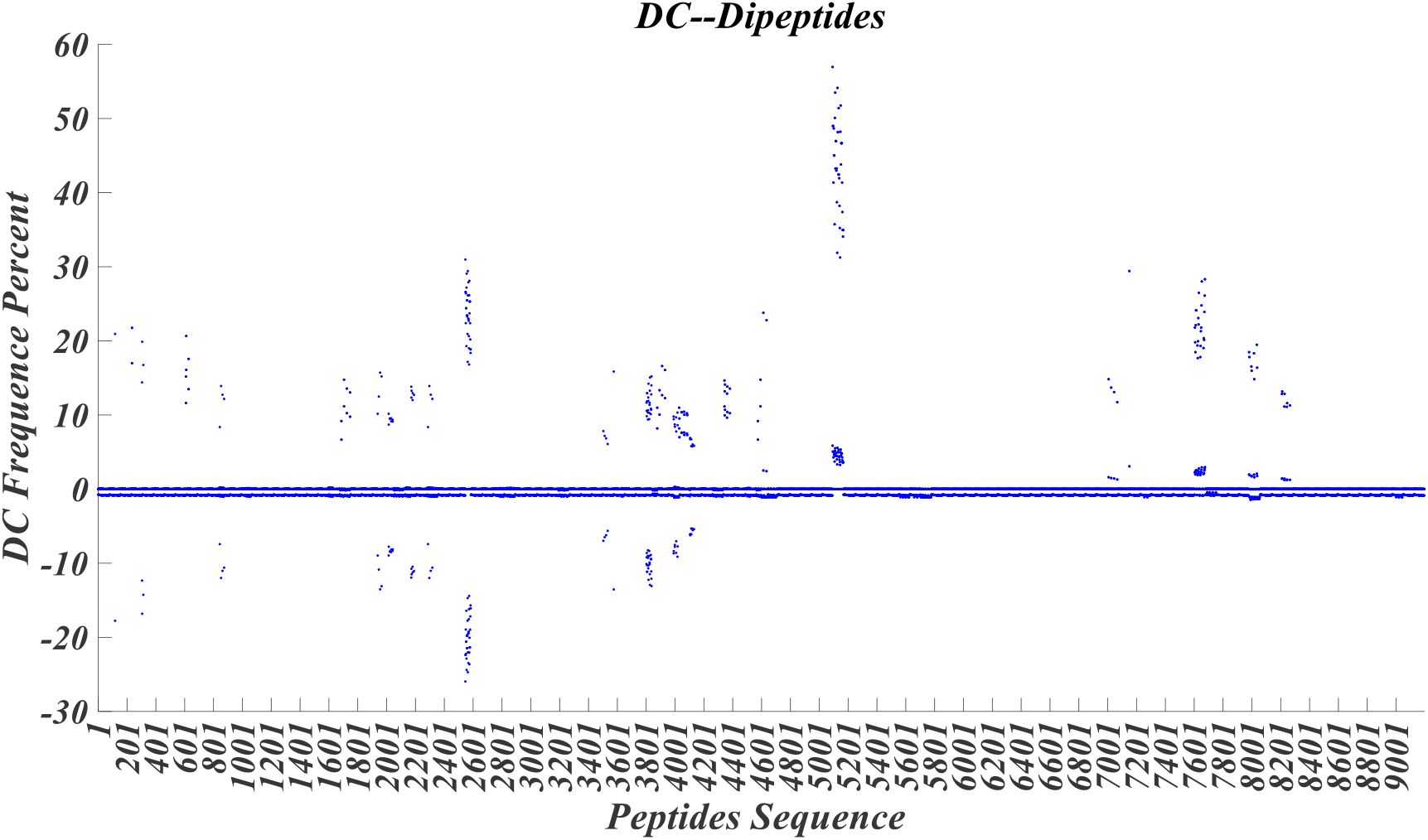
Scatter distribution plots of each peptides sequence

**Fig. 5.**
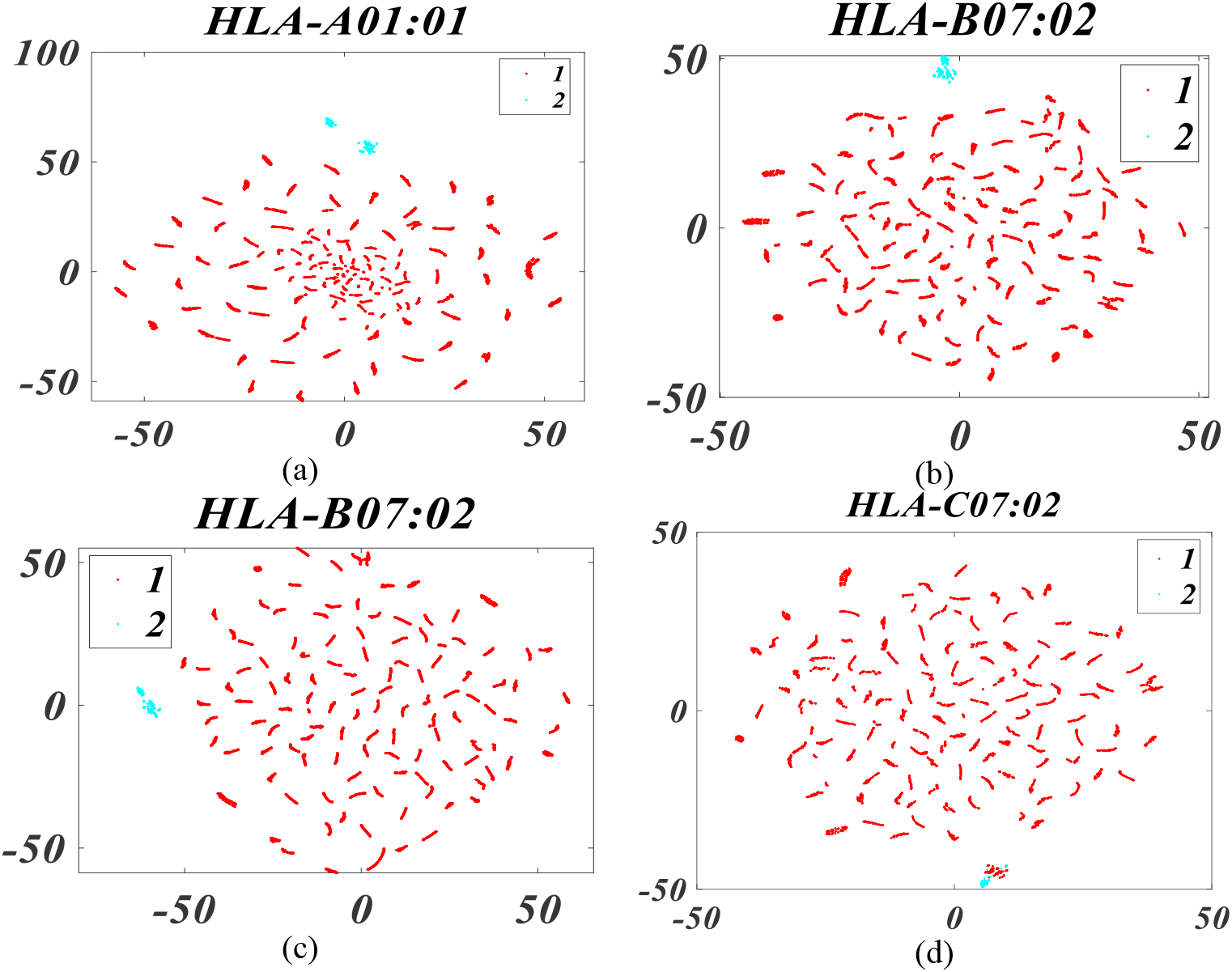
Scatter plots of t-SNE reduced features for the 4 HLA-peptides family. Legend: 1 represents the low DC Frequency & Score of sequences. 2 represents the high DC Frequency & Score of the sequence.

**Fig. 6.**
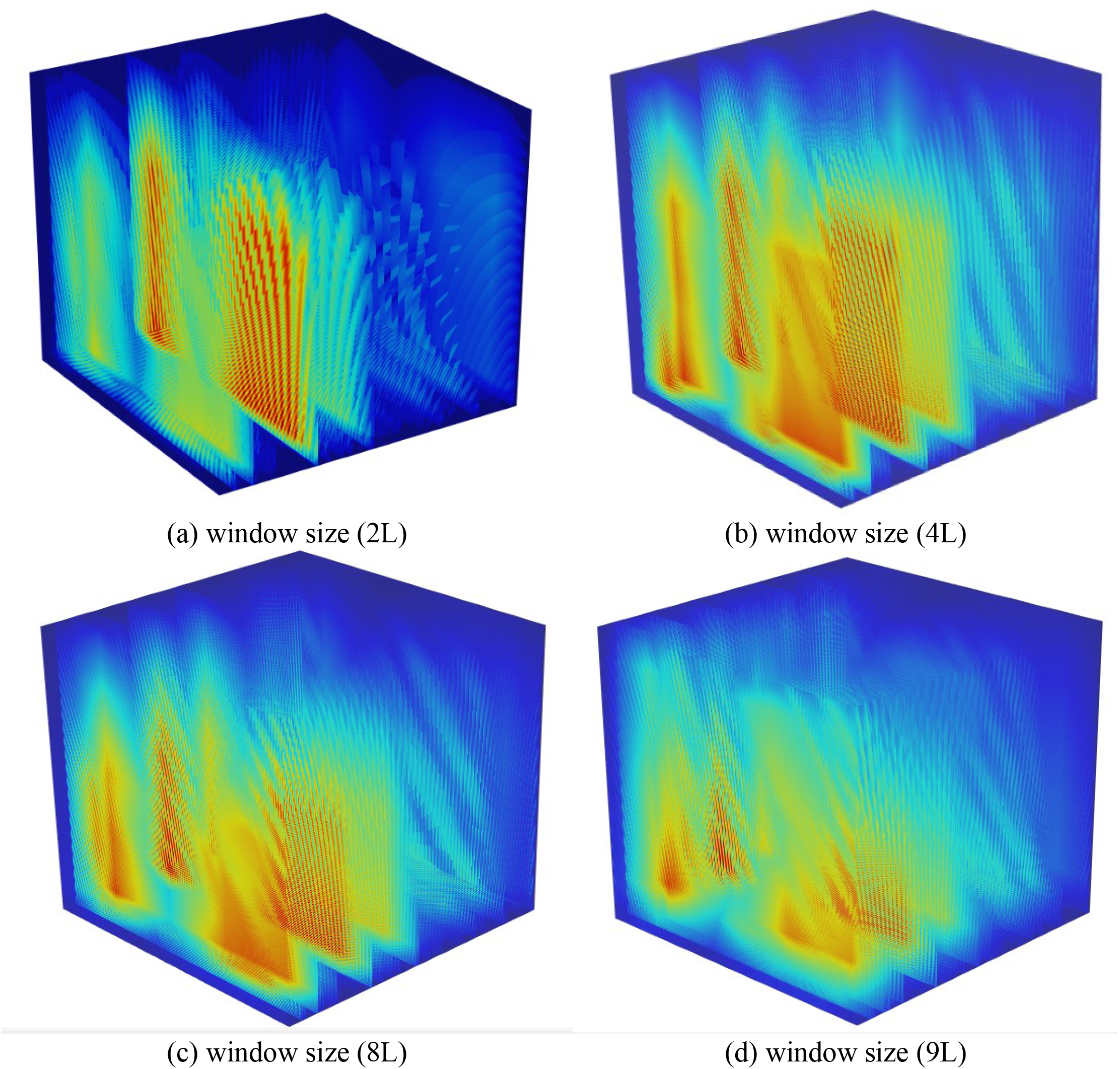
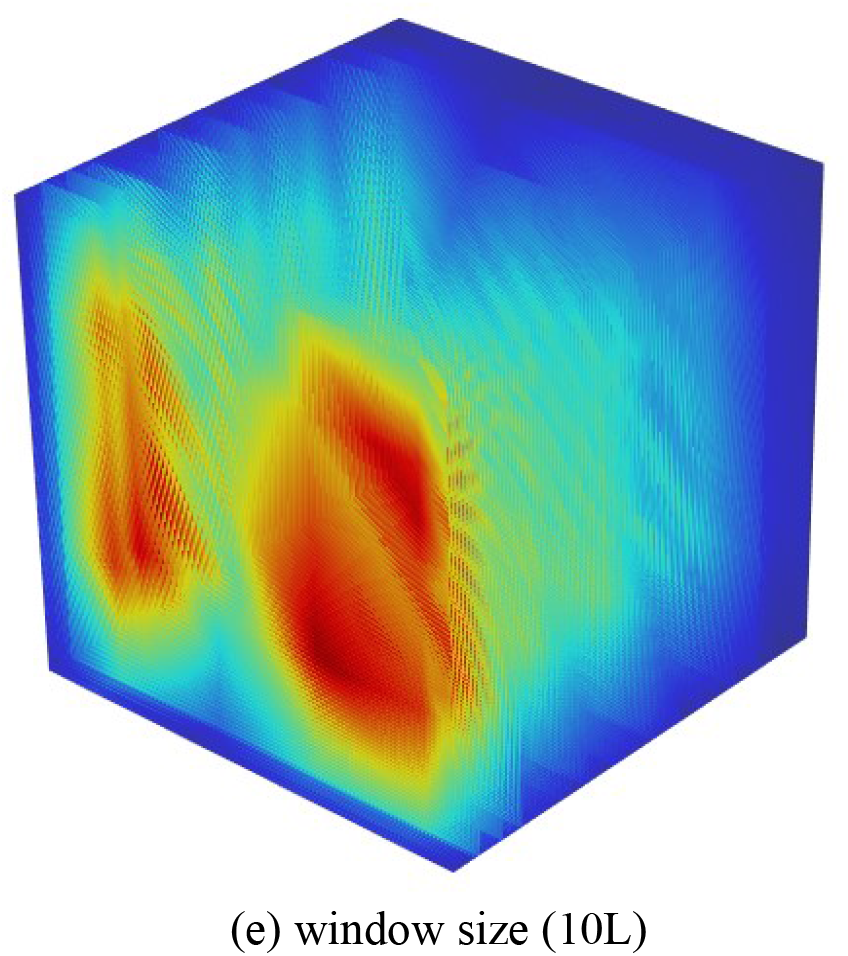
Heatmap Illustrator of each window size (short axis: number of segments across the range of window sizes. Long axis: 382 dipeptides group; the z axis: the HLA peptides dataset from bottom to up)

### 3.1 The Amino Acid Composition (AAC) strategy

AAC[7] computes frequency of all 20 natural amino acid type in a peptide sequence. The fractions of all 20 natural amino acids were calculated as follows followed as:

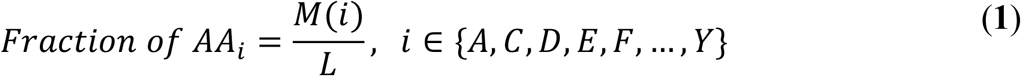

where *M*(*i*) is the total number of specific type *i* of amino acid (AA), while L is the length of the peptide.

It provides numerical vectors of 20 components, with each reflecting the occurrence frequency for the 20 amino acids (sequence order information); This method was developed by [8, 9]to formulate an amino acid sequence of arbitrary length, such as a digital vector. A peptide sequence with length L amino acid residues(Position-Specific-Scoring-Matrix) Sequence composition. For each peptide chain, Equation (1) is used to generate a 1 ×20 dimensional vector according to the amino acid classification.

### 3.2 Local amino acid composition (LAAC)

Amino acid Composition is the normalized frequency of occurrence of each of the twenty amino acids in the given peptide amino acid sequence. Therefore, this feature set includes 20 features[10]. Local enrichment of a single amino acid can dramatically influence the physicochemical properties of a given protein domain, as expressed below:

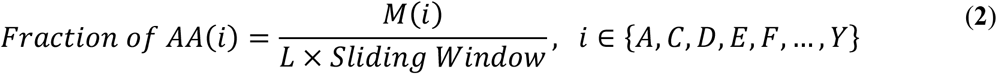

Where Sliding Window =0.1, 0.2, 0.4, 0.6 or 0.8, and L is the length of the peptide., while M(i) is the total number of specific type i of amino acid (AA)

Feature encoding schemes contain the local composition value (per window size) for each amino acid for each peptide and a window size of 5 the weights will be: 0.1aa, 0.2aa, 0.4aa, 0.6aa, 0.8aa. The percent composition of each amino acid is calculated for each window, The lighter the color, the larger the weight of the position. Here, a major goal of our model for binding affinity prediction is to extract insights of the binding mechanism by taking advantage of the attention mechanism, which can learn the importance or contribution of each peptide position to the final binding affinity prediction performance. if the learned neural network pays relatively more attention to some specific amino acid locations of the peptide, these locations may correspond to the binding core. The plot (c), (g), (k), (n), (m), (q), (s) shows an overall significant different.

### 3.3 Dipeptide Composition (DC/DPC)

Since DC can reflect amino acid related information from spatial structure of the sequence[11], we also use dipeptide composition to extract the peptides information DC/DPC provides 382 feature vectors and it can be computed as follows:

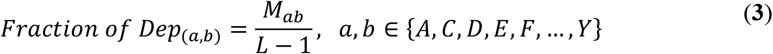

where *M*_*ab*_ is the total number of dipeptides denoted by amino acid types a and b. where dep(a,b) is one dipeptide ab of 382 dipeptides.

Here we display four predictive events (HLA-A01:01; HLA-A03:01; HLA-B07:02; HLA-C07:02 and their corresponding DC frequence percent output vectors. Observations with missing (DC value close to 0) or inconsistent (DC values below 0) information related to the HLA-peptides binding DC information (peptides ID or peptide position on the protein) were removed. We can see that some DC values below 0, the problem of peptide length variety may be one of the causes, which is short peptides length in the dataset.

Furthermore, we have used t-distributed stochastic neighbor embedding (t-SNE) to visualize the features extracted by the original input encoding. Each element represents the weight of the SENnet output. The blue one is the larger the weight of the DC percent in each sub dataset. It suggested that the closer to the potential phosphorylation site, the more important the residue of that position is for the prediction purpose. This finding may provide insights into rules for extracting effective HLA-peptides binding information.

To reduce the redundancy of data and avoid the influence of homology bias, for each DC statistic feature, we sort the values from low to high and use the location to replace the original values. We then divide the orders into 2 groups. The group 1 near the group 2 is strong overlap in the enriched LAAC, which meets our expectation that the predictive events closer to the high LAAC are more important for our model to make predictions.

### 3.4 Local dipeptide composition (LDC)

Compared with other feature expression, less researchers used LDC to study peptides property due to its high dimensionality. In view of this, this paper tries to do some distinctive work with it. Dipeptide Composition [12] describes the proportion of each common amino acid pair within a sequence. It gives 382 features.

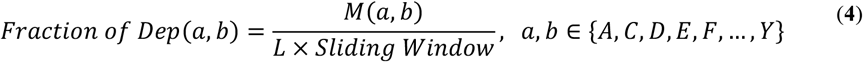

where *M*(*a,b*) is the total number of dipeptides denoted by amino acid types *a* and *b*. where dep(*a,b*) is one dipeptide ab of 382 dipeptides. Window size for 2 4 6 8 9

We utilized with a window’s length as 1, 2, 4, 6 and 8 residues sequence identity cut-off and obtained (10*9196+4*9196+7*9196+) peptide chains. As can be seen, our model correctly identified the 382 dipeptides in the each sliding window size and the none-LDC value (colored blue) or low-LDC value(colored green or light blue). Additionally, most of the high LDC value (colored red or yellow) were also successfully distinguished. Each layer (height) is corresponding to the different HLA sub dataset, and each HLA-peptide binding will be sliced into fragments of 382 dipeptides. the slices in the x-axis direction (long side) respectively represent 382 dipeptides frame, while the z-axis and the description of the scoring metrics used to identify the useful dipeptides based on LDC feature with identical epitopes. also has a strongly negative contribution including NA missing value is screened out, Together, these observations indicate that HLA-pepties binding may tolerate or prefer proteins with specific types of dipeptides including ‘MK’, ‘KR’, ‘RF’, ‘FV’,’VQ’, ‘KH’, ‘HQ’, ‘QK’.

## 4 Network Architecture approaches for predicting NeoAGs

The proposed neural network architecture is shown in Fig. 7. All these sequence data embedding layer types require an initial representation of the elements of the sequence: an initial encoding of the amino acids for target feature vectors.

**Fig. 7.**
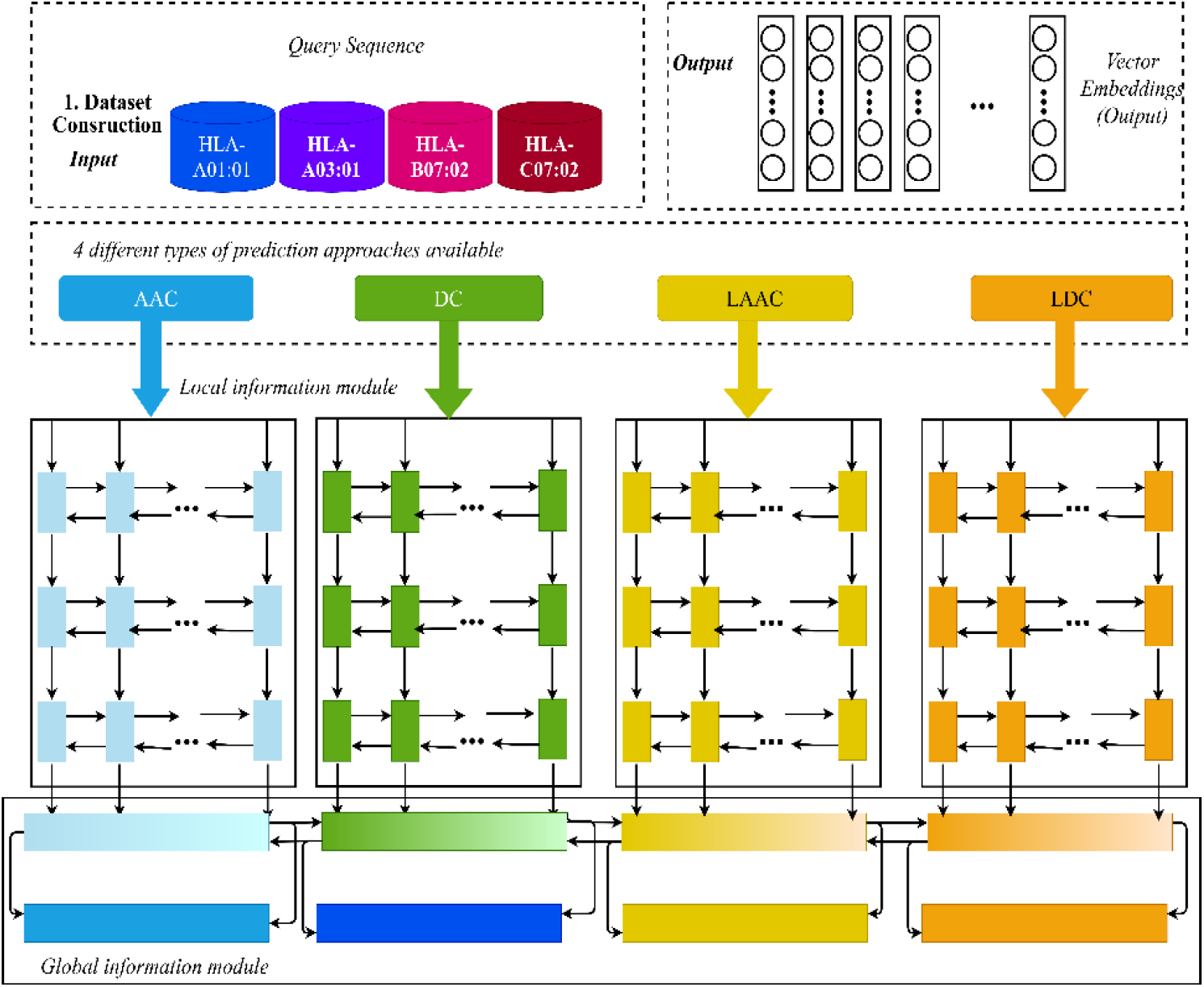
Model Architecture: Brief Overview of DeepNeoAGnet

In our recurrent neural network module, raw peptide and HLA sequences are directly encoded as three vectors of unified sizes. a robust framework for HLA-peptide binding prediction integrating bidirectional LSTM. As a deep learning-based predictor, equipped with a robust network and carefully screened input features, our network’s input carried 4 types of features generated from primary sequences, amino acid features, and profile features. using emerging experimental and computational methods, We set the sliding and put corresponding units in each BiLSTM layer according to the hyperparameter tuning results in this study and merging approaches to predict.

Here, we provide further explanation on the local information processing module. Similar explanation is applicable to the global information processing module because the two modules share the same architecture. The input of the proposed method consisted of both global and local information. In this study, global information referred to the entire peptides sequence information, which included the peptides sequence and peptides phosphorylation feature value by fusing multiple feature representation; while the peptide sequence or any the possible phosphorylation feature value was separately treated as local information.

Finally, the local information-processing module produced an output called as local features. Note that all features need to be fed to the corresponding full connection layer to produce more advanced features. For the global information-processing module, the global features can be obtained using a similar architecture.

### 4.1 Input layer

The encoder takes a sequence of amino acids representing a HLA-peptide binding and encodes it into a sequence of vector representations of the same length. To allow the vector representations at each position to be functions of all surrounding amino acids, we structure the encoder as a stack of bidirectional LSTMs followed by a linear layer projecting the outputs of the last biLSTM layer into the final embedding space.

Input layer transfers sequence region among sliding windows to (20+20×(2+4+7+9+10)+382+382×(2+4+7+9+10)) × 9196 matrices. These representing matrices will then be fed into local information module. Each node in local information module corresponds to one residue.

### 4.2 DeepNeoAGNet local information module layer

In our DeepNeoAGNet model, local information module layer can be seen as feature extractors for HLA-peptides binding residues’ local environment. Pattern extracted by this layer will be passed to global information module for final prediction. As shown in Fig. 7, local information module is a many-to-one bilstm neural network. Local information module for each sliding centered window consists of two directional LSTM layers (forward and backward), each LSTM layer has corresponding memory blocks, corresponds to feature number residues in each centered window. In our study, local information module only output the hidden layer state of the last time step on each direction. Then forward and backward hidden state are concatenated. After processing, local information module will generate one hidden state to represent each centered window.

### 4.3 DeepNeoAGNet global information module layer

The global information module layer, features extracted from the same HLA-peptides binding by local information module will be analyzed together in one bilstm network. As shown in Fig. 7, global information module is a many-to-many bilstm neural network. For each sequence position, global information module will return a concatenated hidden state. In this way, the inter-residue relationships will be effectively learned and facilitate the prediction for every residue by considering the mutual effects between different peptides sequence properties.

### 4.4 Sequence distributed output layer

Sequence distributed layer is a wrapper, which conduct output for every node in global information module layer. Each node in output layer will generate a binary vector to demonstrate the probabilities of bonding states. The binary vector consists of the probabilities of two bonding statements (bonded or free) for the corresponding residue. The statement with higher probability will be selected as the prediction. This prediction is based on the hidden state of the corresponding global information module sequence location.

## 5 The AG prediction tools QA Quality assessment

For evaluation, 90% of the data are used for training, and 10% for testing. We evaluated the performance of the full model, as well as other models trained on each possible combination of biological features. To fully evaluate the performance of DeepNeo-ABNet and related methods, we use the average RMS and R-square values as the final prediction scores. Evaluating of immunogenicity of candidate neoantigens, reliable performances have been achieved regarding this method, which refer to results of the proposed predictor and the state-of-the-art predictors on 4 HLA independent dataset.

### 5.1 RMS (Root-Mean-Square Level)

The calculation formulas of these evaluation parameters were illustrated as follows:

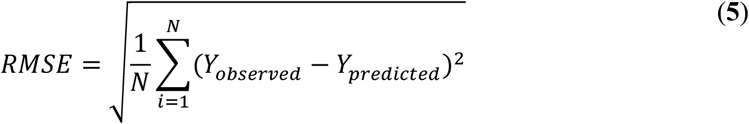

where N is the size of train set or test set, Y*predicted* is the predicted value of in sequence *i* at each peptide epitope, and Y*observed* is the corresponding ground truth.

### 5.2 R Squared (Efficient of Determination)

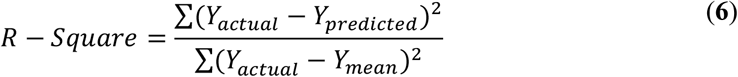

Analyzing cancer neoantigen candidates with numerical series, the positional conservation of amino acids in HLA-A01:01, HLA-A03:01, HLA-B07:02 and HLA-C07:02 is examined by using a analysis, as shown in Fig. 8. Peptides of various lengths need to be proceeded with extra processes to have a fix-length sequence.

**Fig. 8.**
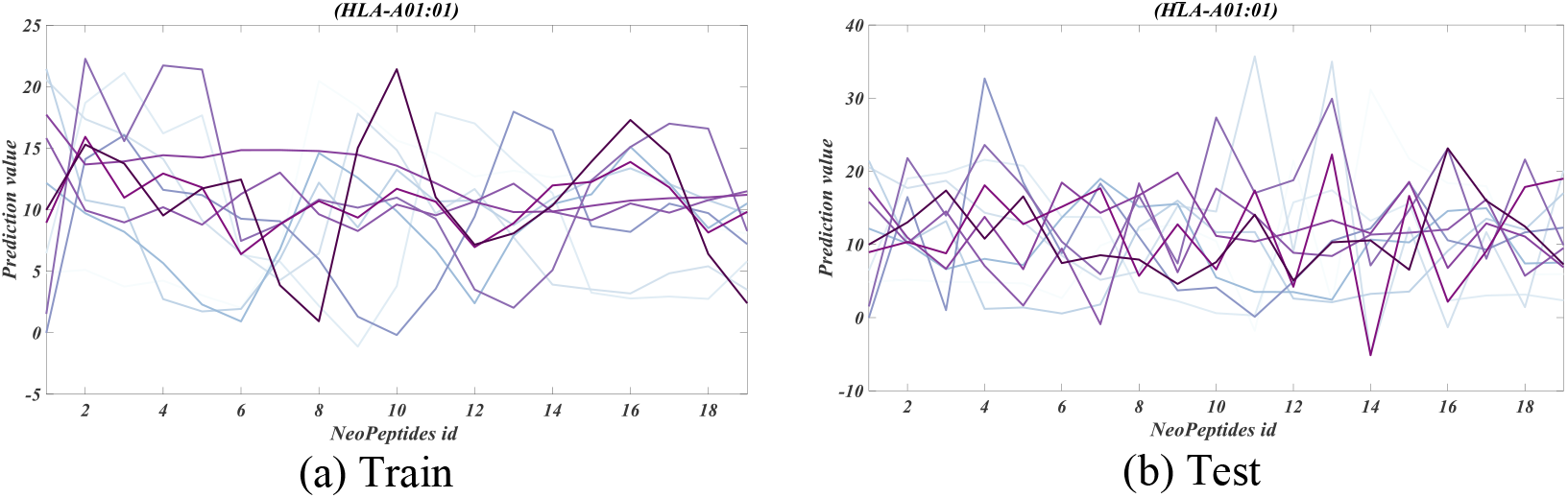
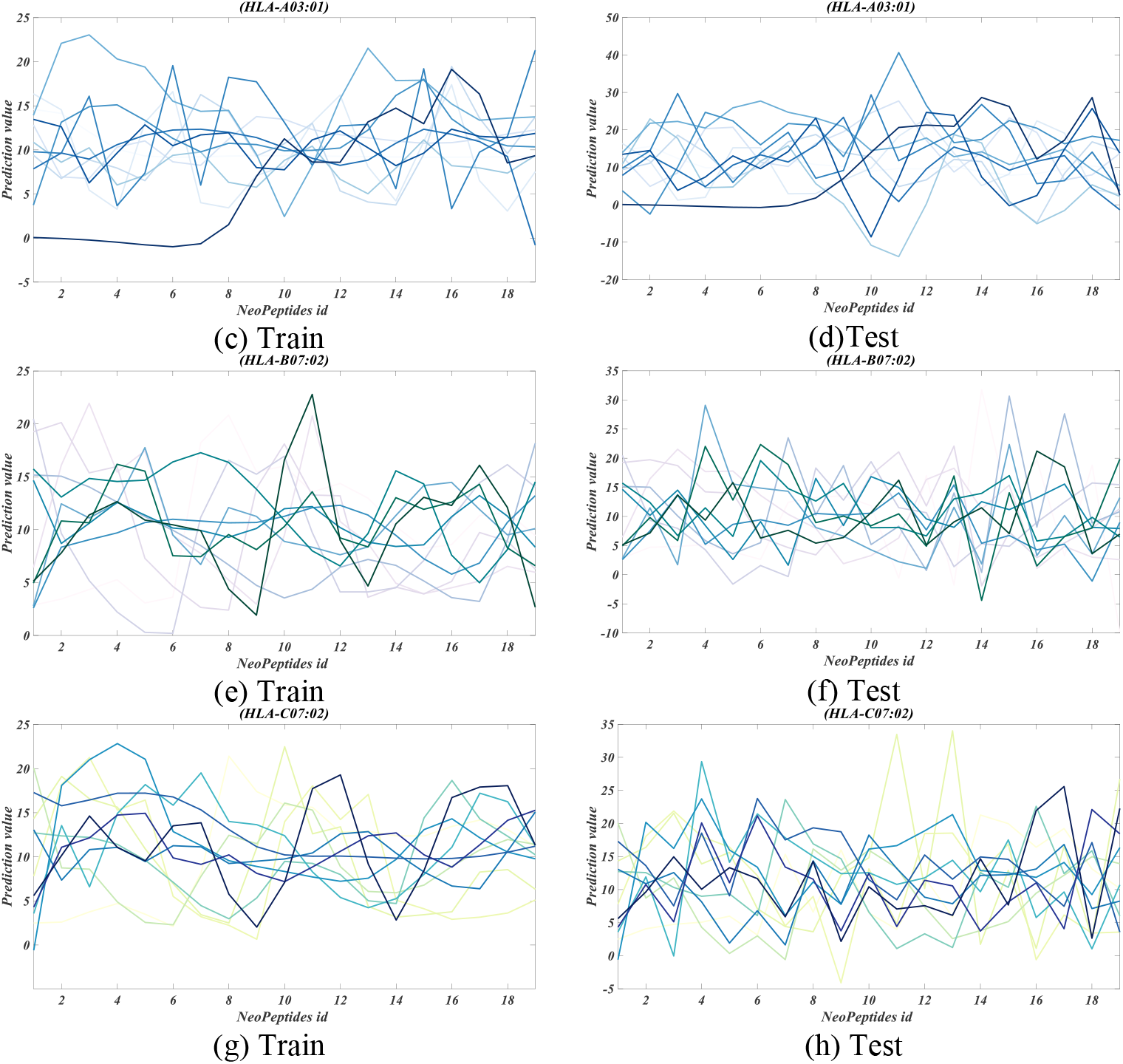
Prediction Results of 4 Different Predictive tasks of DeepNeoAGNet on Train and Test Dataset

**Fig. 9.**
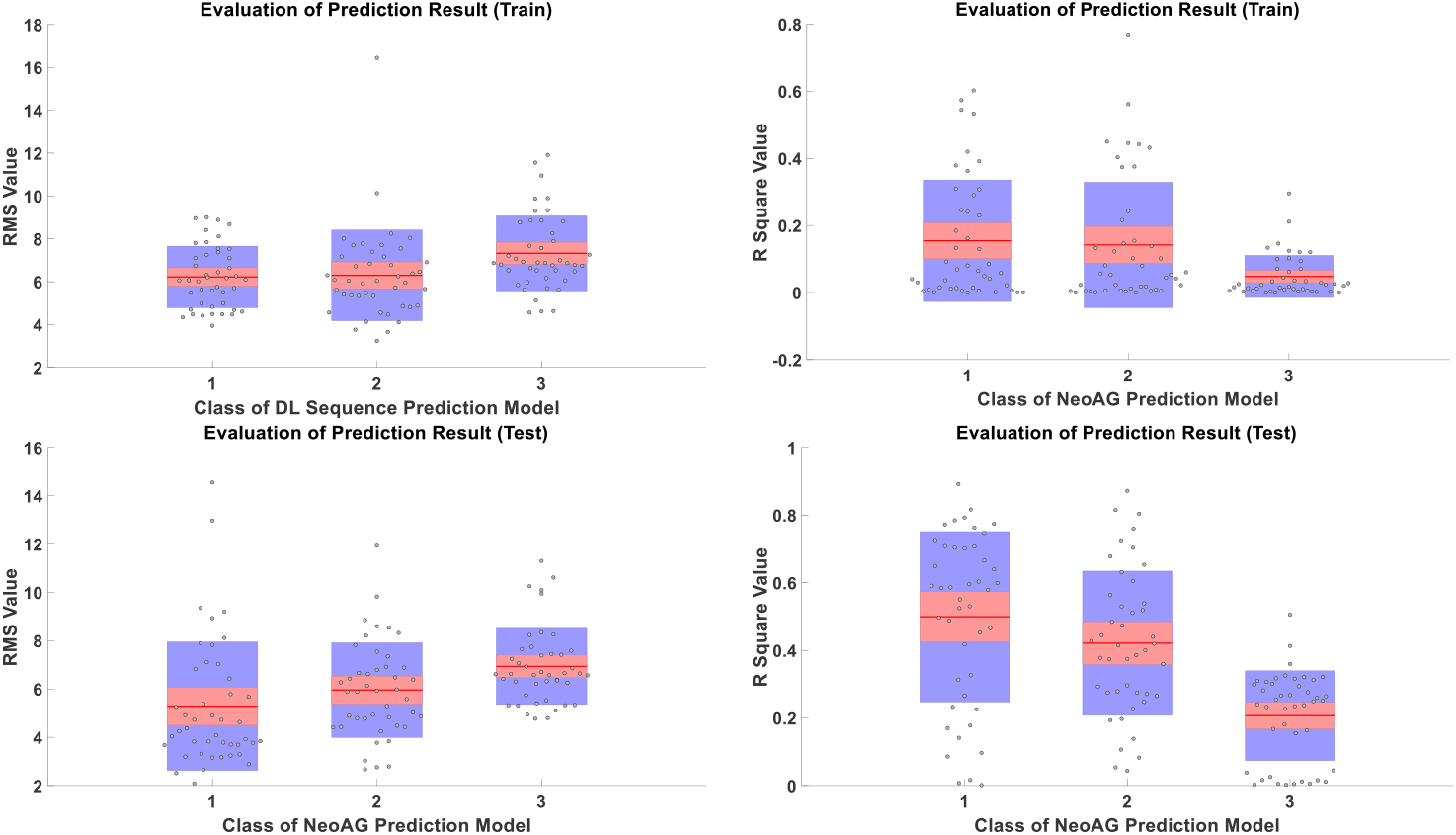
Overall Peptides Binding Prediction Effect to train and test. 1,2,3, is different prediction methods based on the independent dataset Comparisons of error rate

**Table 1.** DeepNeoAGNet Result based on 4 training dataset

| Train |  |  |  |  |  |  |  |  |  |  |  |  |
| --- | --- | --- | --- | --- | --- | --- | --- | --- | --- | --- | --- | --- |
|  | Evaluation index | Seq_1 | Seq_2 | Seq_3 | Seq_4 | Seq_5 | Seq_6 | Seq_7 | Seq_8 | Seq_9 | Seq_10 | Seq_11 |
| **RMS** | HLA-A01:01 | **6.7480** | 7.8100 | 5.4860 | 6.0654 | 8.9655 | **6.3192** | 4.6957 | 4.9854 | 6.0691 | 6.0162 | 7.1045 |
|  | HLA-A03:01 | 9.0096 | **4.3359** | 6.2049 | 5.7539 | **5.6372** | 8.8857 | 6.1679 | 7.2557 | 5.5879 | 4.8263 | **3.9445** |
|  | HLA-B07:02 | 8.4162 | 6.2557 | **4.4733** | **4.4172** | 7.3751 | 7.0997 | **4.4909** | **4.4793** | **5.5086** | 4.9969 | 6.4400 |
|  | HLA-C07:02 | 8.6809 | 8.1149 | 5.6995 | 7.8480 | 7.5464 | 6.6344 | 4.4697 | 4.6020 | 7.5375 | **4.6777** | 6.1009 |
| **R_square** | HLA-A01:01 | **0.1334** | 0.0923 | 0.3095 | 0.2459 | 0.1843 | 0.0400 | 0.3787 | 0.0301 | 0.0045 | 0.0087 | 0.0008 |
|  | HLA-A03:01 | 0.0156 | **0.5731** | 0.0365 | 0.2412 | **0.2900** | 0.0115 | 0.0688 | **0.0803** | 0.0132 | 0.0642 | **0.6022** |
|  | HLA-B07:02 | 0.0045 | 0.2293 | **0.5442** | **0.5330** | 0.0000 | 0.0139 | 0.3622 | 0.0061 | **0.4196** | 0.0489 | 0.0407 |
|  | HLA-C07:02 | 0.0013 | 0.0580 | 0.3074 | 0.0216 | 0.0066 | **0.0847** | **0.3912** | 0.0008 | 0.0006 | **0.1614** | 0.1299 |

Note: Best performing method in **bold**.

**Table 2.** DeepNeoAGNet Result

|  |  | Test |  |  |  |  |  |  |  |  |  |  |
| --- | --- | --- | --- | --- | --- | --- | --- | --- | --- | --- | --- | --- |
|  | Evaluation in-<br>dex | Seq_1 | Seq_2 | Seq_3 | Seq_4 | Seq_5 | Seq_6 | Seq_7 | Seq_8 | Seq_9 | Seq_10 | Seq_11 |
| RMS | HLA-A01:01 | **4.4117** | 5.8838 | 7.8215 | **3.0315** | 5.8815 | **6.2741** | 6.1293 | **2.6719** | **4.4281** | **4.8939** | 4.7943 |
|  | HLA-A03:01 | 6.4286 | 4.7717 | 8.2140 | 6.6607 | 6.6162 | 9.8249 | 11.9325 | 8.8542 | 6.7899 | 7.5433 | 4.9306 |
|  | HLA-B07:02 | 7.3461 | **2.7608** | **4.2508** | 5.9263 | 8.5997 | 8.3230 | 6.9187 | 2.7921 | 4.8324 | 5.2908 | **4.4788** |
|  | HLA-C07:02 | 6.4771 | 5.9783 | 5.5807 | 3.7715 | **4.4277** | 6.8819 | **5.0306** | 3.8361 | 8.5345 | 6.3902 | 4.8681 |
| R_square | HLA-A01:01 | **0.5633** | 0.4279 | 0.3778 | **0.8150** | 0.2929 | 0.4450 | 0.4844 | **0.6778** | 0.3733 | **0.4730** | 0.1933 |
|  | HLA-A03:01 | 0.4145 | 0.5298 | 0.2751 | 0.7255 | 0.3747 | 0.0538 | 0.1062 | 0.1970 | 0.0437 | 0.0827 | **0.6313** |
|  | HLA-B07:02 | 0.2266 | **0.8717** | 0.7034 | 0.5106 | 0.2474 | 0.2787 | 0.2960 | 0.6532 | **0.6055** | 0.3860 | 0.2746 |
|  | HLA-C07:02 | 0.2710 | 0.4003 | **0.7597** | 0.8040 | **0.5187** | **0.4195** | **0.5388** | 0.4402 | 0.1384 | 0.3598 | 0.2649 |

Note: Best performing method in **bold**.

**Table 3.**
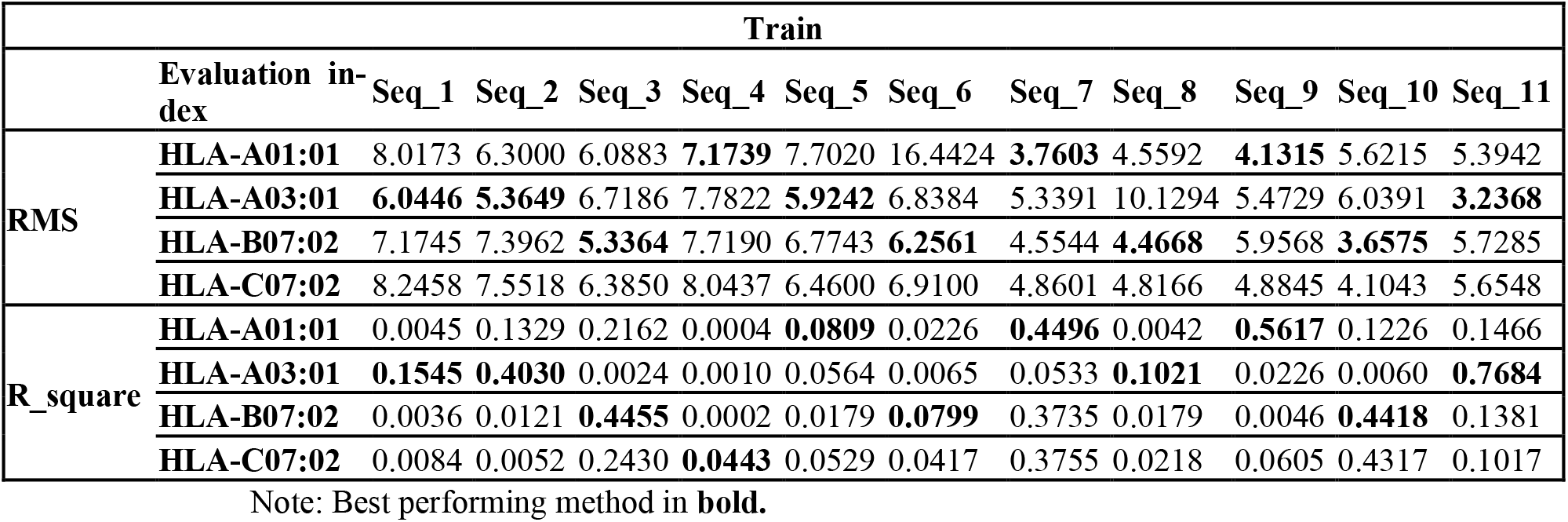
RGU evaluation result

|  |  | Train |  |  |  |  |  |  |  |  |  |  |
| --- | --- | --- | --- | --- | --- | --- | --- | --- | --- | --- | --- | --- |
|  | Evaluation in-<br>dex | Seq_1 | Seq_2 | Seq_3 | Seq_4 | Seq_5 | Seq_6 | Seq_7 | Seq_8 | Seq_9 | Seq_10 | Seq_11 |
| RMS | HLA-A01:01 | 8.0173 | 6.3000 | 6.0883 | **7.1739** | 7.7020 | 16.4424 | **3.7603** | 4.5592 | **4.1315** | 5.6215 | 5.3942 |
|  | HLA-A03:01 | **6.0446** | **5.3649** | 6.7186 | 7.7822 | **5.9242** | 6.8384 | 5.3391 | 10.1294 | 5.4729 | 6.0391 | **3.2368** |
|  | HLA-B07:02 | 7.1745 | 7.3962 | **5.3364** | 7.7190 | 6.7743 | **6.2561** | 4.5544 | **4.4668** | 5.9568 | **3.6575** | 5.7285 |
|  | HLA-C07:02 | 8.2458 | 7.5518 | 6.3850 | 8.0437 | 6.4600 | 6.9100 | 4.8601 | 4.8166 | 4.8845 | 4.1043 | 5.6548 |
| R_square | HLA-A01:01 | 0.0045 | 0.1329 | 0.2162 | 0.0004 | **0.0809** | 0.0226 | **0.4496** | 0.0042 | **0.5617** | 0.1226 | 0.1466 |
|  | HLA-A03:01 | **0.1545** | **0.4030** | 0.0024 | 0.0010 | 0.0564 | 0.0065 | 0.0533 | **0.1021** | 0.0226 | 0.0060 | **0.7684** |
|  | HLA-B07:02 | 0.0036 | 0.0121 | **0.4455** | 0.0002 | 0.0179 | **0.0799** | 0.3735 | 0.0179 | 0.0046 | **0.4418** | 0.1381 |
|  | HLA-C07:02 | 0.0084 | 0.0052 | 0.2430 | **0.0443** | 0.0529 | 0.0417 | 0.3755 | 0.0218 | 0.0605 | 0.4317 | 0.1017 |

Note: Best performing method in **bold**.

**Table 4.** RGU evaluation result

|  |  | Test |  |  |  |  |  |  |  |  |  |  |
| --- | --- | --- | --- | --- | --- | --- | --- | --- | --- | --- | --- | --- |
|  | Evaluation in-<br>dex | Seq_1 | Seq_2 | Seq_3 | Seq_4 | Seq_5 | Seq_6 | Seq_7 | Seq_8 | Seq_9 | Seq_10 | Seq_11 |
| RMS | HLA-A01:01 | **3.6769** | **2.5159** | 5.2580 | 3.1980 | 9.3568 | 4.9164 | **4.0306** | **2.0843** | **2.6552** | 4.2492 | 6.8331 |
|  | HLA-A03:01 | 8.9326 | 4.7263 | 5.3797 | 7.8983 | **4.3678** | 14.5503 | 12.9738 | 7.1076 | 9.2031 | 7.0339 | **4.8987** |
|  | HLA-B07:02 | 3.8252 | 3.3186 | **3.8249** | **3.1578** | 7.8286 | 4.7235 | 4.0836 | 3.1706 | 3.2440 | **3.2926** | 8.1149 |
|  | HLA-C07:02 | 3.7823 | 3.7022 | 6.4307 | 3.6886 | 5.7856 | **3.9225** | 4.6336 | 3.7705 | 2.8888 | 3.8408 | 5.6743 |
| R_square | HLA-A01:01 | 0.7263 | **0.8918** | 0.6494 | **0.7721** | 0.2328 | 0.5912 | **0.5842** | **0.7845** | **0.7081** | 0.5862 | 0.1699 |
|  | HLA-A03:01 | 0.2656 | 0.5505 | 0.4175 | 0.3131 | **0.4967** | 0.0861 | 0.1412 | 0.0073 | 0.0165 | 0.0015 | **0.5954** |
|  | HLA-B07:02 | 0.7036 | 0.8164 | **0.7927** | 0.7628 | 0.3264 | 0.4881 | 0.5248 | 0.6034 | 0.5786 | **0.7013** | 0.1775 |
|  | HLA-C07:02 | **0.7471** | 0.7742 | 0.5304 | 0.7070 | 0.0967 | **0.6658** | 0.4530 | 0.4659 | 0.6397 | 0.5993 | 0.2258 |

Note: Best performing method in **bold**.

**Table 5.**
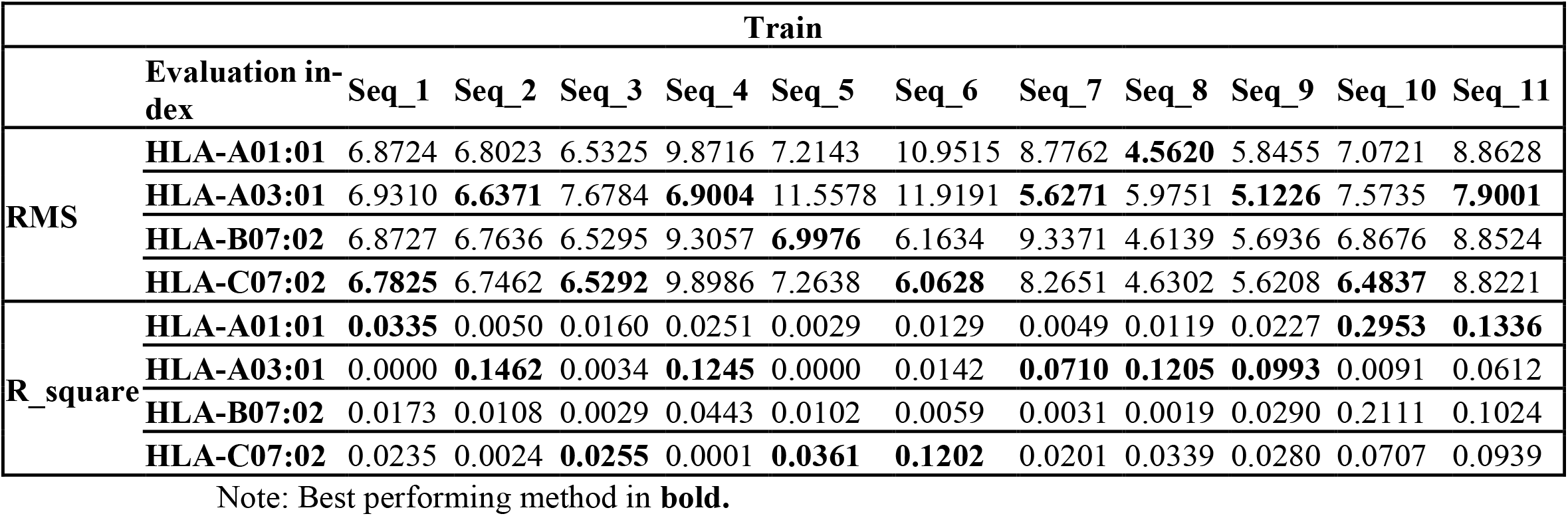
BILSTM evaluation result

**Table 6.**
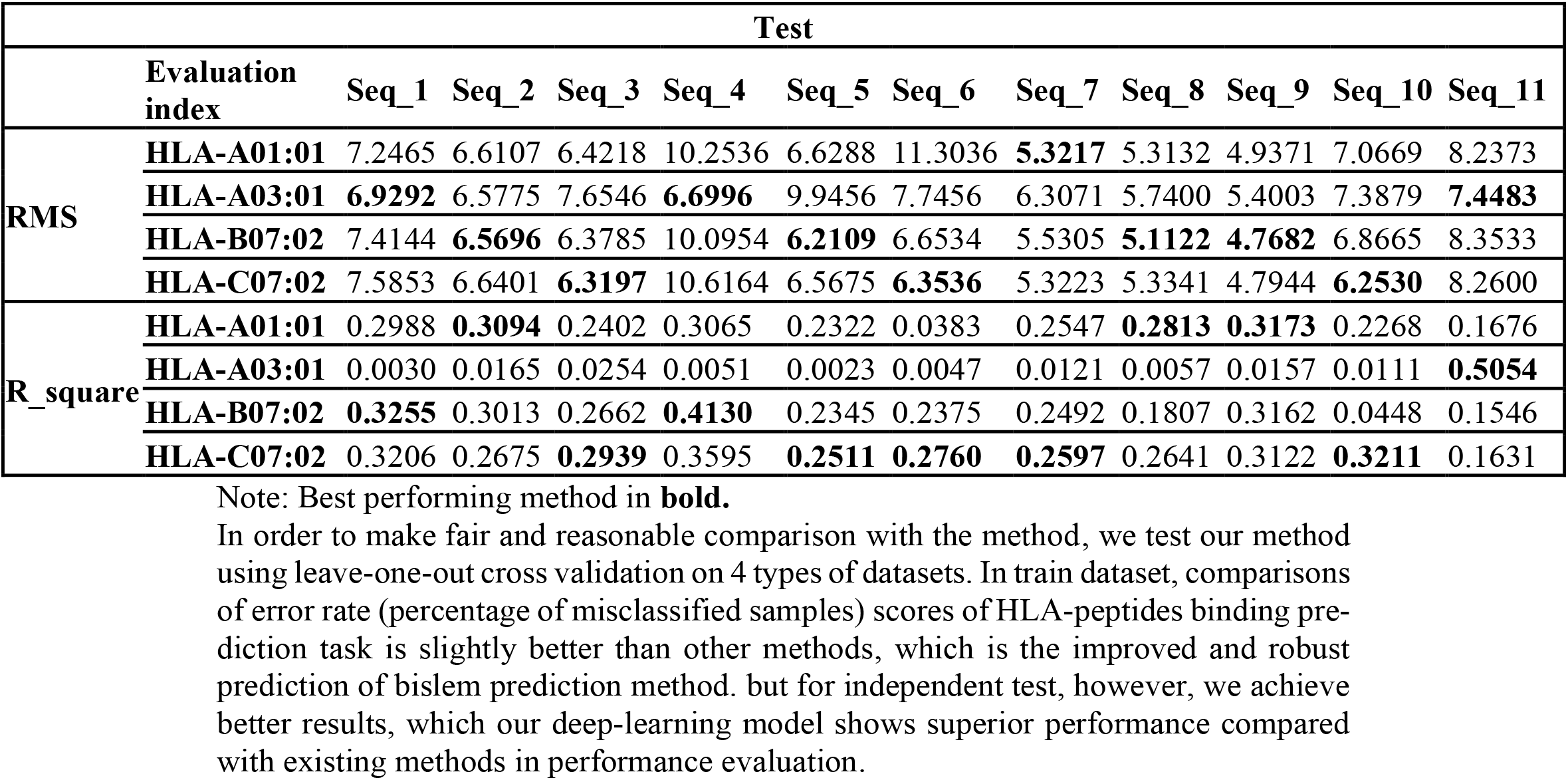
BILSTM evaluation result

|  |  | Test |  |  |  |  |  |  |  |  |  |  |
| --- | --- | --- | --- | --- | --- | --- | --- | --- | --- | --- | --- | --- |
|  | Evaluation index | Seq_1 | Seq_2 | Seq_3 | Seq_4 | Seq_5 | Seq_6 | Seq_7 | Seq_8 | Seq_9 | Seq_10 | Seq_11 |
| RMS | HLA-A01:01 | 7.2465 | 6.6107 | 6.4218 | 10.2536 | 6.6288 | 11.3036 | **5.3217** | 5.3132 | 4.9371 | 7.0669 | 8.2373 |
|  | HLA-A03:01 | **6.9292** | 6.5775 | 7.6546 | **6.6996** | 9.9456 | 7.7456 | 6.3071 | 5.7400 | 5.4003 | 7.3879 | **7.4483** |
|  | HLA-B07:02 | 7.4144 | **6.5696** | 6.3785 | 10.0954 | **6.2109** | 6.6534 | 5.5305 | **5.1122** | **4.7682** | 6.8665 | 8.3533 |
|  | HLA-C07:02 | 7.5853 | 6.6401 | **6.3197** | 10.6164 | 6.5675 | **6.3536** | 5.3223 | 5.3341 | 4.7944 | **6.2530** | 8.2600 |
| R_square | HLA-A01:01 | 0.2988 | **0.3094** | 0.2402 | 0.3065 | 0.2322 | 0.0383 | 0.2547 | **0.2813** | **0.3173** | 0.2268 | 0.1676 |
|  | HLA-A03:01 | 0.0030 | 0.0165 | 0.0254 | 0.0051 | 0.0023 | 0.0047 | 0.0121 | 0.0057 | 0.0157 | 0.0111 | **0.5054** |
|  | HLA-B07:02 | **0.3255** | 0.3013 | 0.2662 | **0.4130** | 0.2345 | 0.2375 | 0.2492 | 0.1807 | 0.3162 | 0.0448 | 0.1546 |
|  | HLA-C07:02 | 0.3206 | 0.2675 | **0.2939** | 0.3595 | **0.2511** | **0.2760** | **0.2597** | 0.2641 | 0.3122 | **0.3211** | 0.1631 |

Note: Best performing method in **bold**.
In order to make fair and reasonable comparison with the method, we test our method using leave-one-out cross validation on 4 types of datasets. In train dataset, comparisons of error rate (percentage of misclassified samples) scores of HLA-peptides binding prediction task is slightly better than other methods, which is the improved and robust prediction of bislem prediction method. but for independent test, however, we achieve better results, which our deep-learning model shows superior performance compared with existing methods in performance evaluation.

## 6 Results and discussion

Our work is shown to reliably predict neoantigen of a given target antigen sequence, thus contributing to a significant reduction of laboratory experiments and costs required for the conventional approach. DeepNeoAGNet outperforms state-of-the-art model by in terms of RMS and R-square respectively. DeepNeoAGNet was user-friendly with no length limitation of input and had the highest output efficiency among the existing methods with an acceptable RMSE and R-square.

## Conclusion and future development

Neoantigens play important roles in cancer immunotherapy. Current methods used for neoantigen prediction focus on the binding between human leukocyte antigens (HLAs) and peptides, which is insufficient for high-confidence neoantigen prediction. In this study, we apply deep learning techniques to predict neoantigens considering both the possibility of HLA-peptide binding and the potential immunogenicity of the peptide-HLA. This method, in association with a similarity search tool, can be used for automated annotation of peptides data. The binding model achieves comparable performance with other well-acknowledged tools on the latest Immune Epitope Database (IEDB) benchmark datasets and an independent mass spectrometry (MS) dataset. The Global-Local module could significantly improve the prediction precision of neoantigens. Understanding and predicting HLA–peptide binding is an essential step for studies of the immune system, T-cell epitopes, and adverse drug reactions. The methods that are popular in the field of computer sciences are widely used for HLA–peptide binding predictions. Different types of descriptors were utilized to transfer the peptide and HLA sequences into model-acceptable numbers. Some extra processes were implemented to deal with peptides with various lengths.

We evaluated rms and r-square on HIA dataset and proved that it achieves comparable performance with the state-of-the-art methods. Our work has been actively refining a machine-learning model to improve neoantigen identification and predictions. Deep-NeoAGNet moel could support the personalized neoantigen-based cancer vaccine and cell therapy development.

However, there is still room for further investigation. For example, the problem of an imbalanced dataset had a negative effect on the accuracy of small-sized types. We just exact the high percent of feature value, and No conclusion was drawn to indicate which one is absolutely better than the other. Despite the impressive advances in using DeepNeoAGNet as potent therapeutics, there are still many challenges that continue to frustrate their efficient development. Besides the physicochemical properties of amino acids, such as hydrogen bond number, polarity, and hydrophobicity were used as descriptors. To capture more contextual information, the LAAC, LDC, GD and LZC methods consider different amino acid classification approaches. Although our method obtains relatively satisfactory results, some open problems need to be investigated in the future.

The further application of our method to the mutations with pre-existing T-cell responses indicating its feasibility in clinical application. In future work, we intend to verify a number of the hypotheses provided here by engineering proteins to better reflect the features found to be related to high production rates. The identification of neoantigens as drivers of successful antitumor immunity is offering exciting new opportunities for cancer immunotherapies, including making T-cell infusion products highly individualized for more effective treatment. We plan to extend the model training on more variable HLA alle peptides in future work.

## Data Availability

All relevant data are within the paper and its Supporting Information files.

## Acknowledgments

N/A, not available/applicable.

## References

1. Abelin, J. G. et al. Mass spectrometry profiling of HLA-associated peptidomes in mono-allelic cells enables more accurate epitope prediction. Immunity 46, 315–326 (2017).

2. Cohen, C. J. et al. Isolation of neoantigen-specific T cells from tumor and peripheral lymphocytes. J. Clin. Invest. 125, 3981–3991 (2015).

3. Liu, Z., Cui, Y., Xiong, Z. et al. DeepSeqPan, a novel deep convolutional neural network model for pan-specific class I HLA-peptide binding affinity prediction. Sci Rep 9, 794 (2019). https://doi.org/10.1038/s41598-018-37214-1

4. Liu Z, Jin J, Cui Y, Xiong Z, Nasiri A, Zhao Y, Hu J. DeepSeqPanII: an interpretable recurrent neural network model with attention mechanism for peptide-HLA class II binding prediction. IEEE/ACM Trans Comput Biol Bioinform. 2021 Apr 22;PP. doi: 10.1109/TCBB.2021.3074927. Epub ahead of print. PMID: 33886473.

5. Gartner, J.J., Parkhurst, M.R., Gros, A. et al. A machine learning model for ranking candidate HLA class I neoantigens based on known neoepitopes from multiple human tumor types. Nat Cancer 2, 563–574 (2021). https://doi.org/10.1038/s43018-021-00197-6

6. Mei S. Multi-kernel transfer learning based on Chou’s PseAAC formulation for protein sub-mitochondria localization. J. Theor. Biol. 2012;293:121–130.

7. K.C. Prediction of protein cellular attributes using pseudo-amino acid composition. Proteins. 2001;43:246–255.

8. X.Wei R., Zhang T., Gu Q. Using the concept of Chou’s pseudo amino acid composition to predict apoptosis proteins subcellular location: An approach by approximate entropy. Protein Pept. Lett. 2008;15:392–396.

9. Lin H., Wang H., Ding H., Chen Y.-L., Li Q.-Z. Prediction of subcellular localization of apoptosis protein using Chou’s pseudo amino acid composition. Acta Biotheor. 2009;57:321–330.

10. Saini SK, Ostermeir K, Ramnarayan VR, Schuster H, Zacharias M, Springer S. Dipeptides promote folding and peptide binding of MHC class I molecules. Proc Natl Acad Sci U S A. 2013 Sep 17;110(38):15383–8. doi: 10.1073/pnas.1308672110. Epub 2013 Sep 3. PMID: 24003162; PMCID: PMC3780906.

11. Zhang, YH., Xing, Z., Liu, C. et al. Identification of the core regulators of the HLA I-peptide binding process. Sci Rep 7, 42768 (2017). https://doi.org/10.1038/srep42768

12. Khan, A. R., Baker, B. M., Ghosh, P., Biddison, W. E. & Wiley, D. C. The structure and stability of an HLA-A*0201/octameric tax peptide complex with an empty conserved peptide-N-terminal binding site. Journal of immunology 164, 6398–6405 (2000).

